# Identifying developing interneurons as a potential target for multiple genetic autism risk factors in human and rodent forebrain

**DOI:** 10.1101/2021.06.03.446920

**Authors:** Yifei Yang, Sam A. Booker, James M. Clegg, Idoia Quintana Urzainqui, Anna Sumera, Zrinko Kozic, Owen Dando, Sandra Martin Lorenzo, Yann Herault, Peter C. Kind, David J. Price, Thomas Pratt

## Abstract

Autism spectrum condition or ‘autism’ is associated with numerous monogenic and polygenic genetic risk factors including the polygenic *16p11.2* microdeletion. A central question is what neural cells are affected. To systematically investigate we analysed single cell transcriptomes from gestational week (GW) 8-26 human foetal prefrontal cortex and identified a subset of interneurons (INs) first appearing at GW23 with enriched expression of a disproportionately large fraction of risk factor transcripts. This suggests the hypothesis that these INs are disproportionately vulnerable to mutations causing autism. We investigated this in a rat model of the *16p11.2* microdeletion. We found no change in the numbers or position of either excitatory or inhibitory neurons in the somatosensory cortex or CA1 of *16p11.2*^*+/-*^ rats but found that CA1 Sst INs were hyperexcitable with an enlarged axon initial segment, which was not the case for CA1 pyramidal cells. This study prompts deeper investigation of IN development as a convergent target for autism genetic risk factors.

## INTRODUCTION

Autism spectrum conditions (ASC – referred to here as ‘autism’) describe several symptoms and behaviours which affect the way in which a group of people understand and react to the world around them (Mental Health Foundation) and may co-occur with other conditions including epilepsy and intellectual disability (ID). Recent efforts to understand the genetic landscape of autism identified hundreds of genetic risk factors predisposing to autism including *de novo* single gene mutation (5-10%), copy number variations (CNVs) and chromosome anomalies (5%), and inherited single gene mutations (3%), although most cases (80-85%) have no known genetic cause.

Genetic risk factors can be either ‘monogenic’ where a single gene mutation is sufficient to predispose to autism or ‘polygenic’ where one mutation directly affects several genes simultaneously. CNVs are an example of the latter where chromosomal microduplication or microdeletion affect the gene dosage of multiple genes. The Simons Foundation Autism Research Initiative (SFARI) curates a list of ∼1000 monogenic genetic risk factors of which 83 are categorised as the highest ranking (Categories 1 ‘high confidence’ and 2 ‘strong candidates’) indicating a robust association between mutation in these genes and autism. SFARI lists 2240 CNVs associated with autism of which *16p11.2* microdeletion and microduplication are among the most frequent accounting for approximately 1% of cases. The *16p11.2* CNV comprises 27 protein coding genes so is a polygenic risk factor although it is currently unknown the extent to which each of the *16p11.2* genes are individual risk factors.

Autism manifests in early infancy and then persists into later life. A number of lines of evidence suggest that events during brain development *in utero* contribute to the subsequent development of symptoms (Packer, 2016). The developing cerebral cortex is comprised of three neuronal cardinal cell classes (progenitors, excitatory neurons, and inhibitory neurons) and three non-neuronal cardinal cell classes (astrocytes, oligodendrocyte precursors, and microglia). One important factor for cerebral cortex function is the balance between glutamatergic, excitatory principal neurons, which originate from progenitors located in the ventricular zone of the cerebral cortex, and inhibitory GABAergic interneurons (INs) which originate from progenitors located in the ganglionic eminences and then migrate into the cortex and integrate within functional circuits of the cortical plate (Hansen et al., 2013; Ma et al., 2013). Changes in the number, position, anatomy, or electrophysiology of inhibitory or excitatory neurons may perturb the excitatory/inhibitory balance (the E/I balance) and is hypothesised to be a convergent mechanism in autism and its co-occuring conditions (Bozzi et al., 2018; Nelson and Valakh, 2015; Puts et al., 2017; Rapanelli et al., 2017; Robertson et al., 2016) (Antoine et al., 2019)

The aim of the current study is to systematically identify types of cell in developing human cerebral cortex that are potentially vulnerable to autism risk factors using a single cell mRNA sequencing (scRNA-seq) dataset acquired from developing human foetal cortex spanning gestational weeks (GW) 8 to 26 (Zhong et al., 2018). While we found that some autism associated transcripts are differentially expressed between the cardinal cell classes our most striking finding was that a subset of differentiating INs first appearing at GW23 exhibited enriched expression of a strikingly high proportion of risk transcripts. This molecular analysis suggests the hypothesis that a large number of monogenic risk factors and the polygenic *16p11.2* microdeletion selectively target IN development resulting in IN phenotypes postnatally that contribute to autism and its comorbid conditions. We support this hypothesis using a *16p11.2* microdeletion rat model where we identified hypersensitive electrophysiology and enlarged axon initial segment (AIS) phenotypes in somatostatin (Sst) expressing hippocampal INs.

## METHODS

### Datasets

Three published cortical transcriptome datasets were used in this study to explore the gene expression pattern of autism-associated genes during cortical development.

The raw gene expression matrix in the scRNA-seq data of human foetal PFC was obtained from the Gene Expression Omnibus (GEO) under the accession number GSE104276, then the data was normalized as the original paper described (Zhong et al., 2018). We used the authors’ original classification of six cardinal cell classes (NPC, ExN, IN, OPC, Astrocyte and Microglia).

The expression matrix of genes in the adult human cortical single nuclei RNA-seq data were downloaded under the accession number of GSE97930 (Lake et al., 2018). In the dataset from Lake et al., only cells that identified as “INs” were used for further analysis. The original eight interneuron clusters were grouped based on the expression pattern of marker genes (In1/2/3 as VIP, In4 as NG, In6 as PV, In7/8 as SST, Figure 2B,C in Lake et al., 2018).

**Figure 1:**
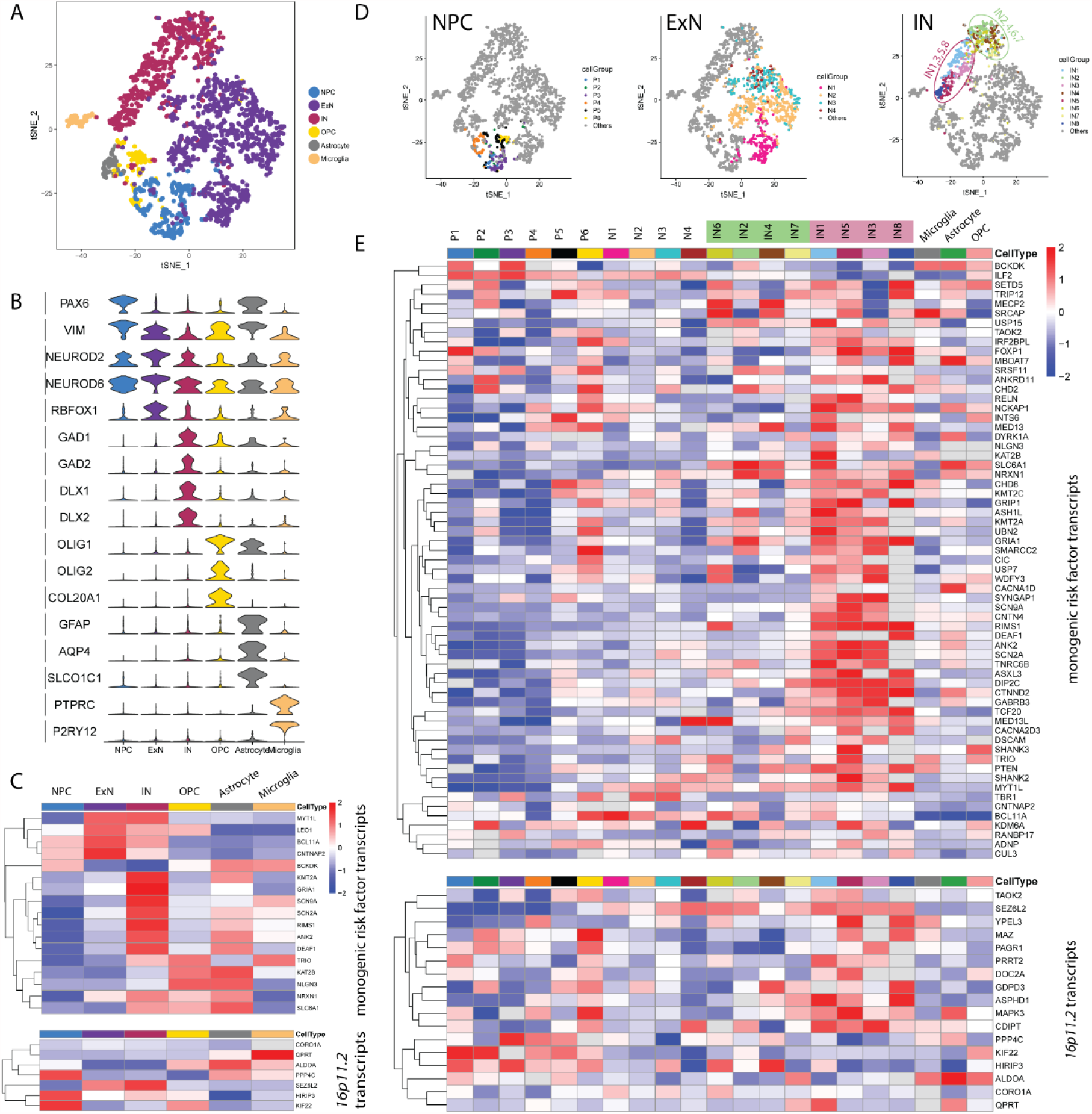
(A-C) Differentially expression of autism risk factor transcripts among foetal cortical cardinal cell classes. (A) *t*-SNE plot showing the cardinal cell classes identified in the dataset. (B) Violin plot illustrating the expression pattern of marker genes among the six cardinal cell classes. (C) Heatmap illustrating the expression pattern of significantly differentially expressed autism risk factor transcripts across cardinal cell classes (Wilcox test, adjust p < 0.05, log (fold change) > 0.3). Top: monogenic autism risk factor transcripts; Bottom: *16p11.2* transcripts. (D,E) Unsupervised clustering within the cardinal classes and similarity comparison between cell clusters. (D) Unsupervised clustering subdividing the cardinal classes into 21 different cell clusters. OPC, astrocytes, and microglia were not further clustered. (E) Heatmap illustrating the expression pattern of differentially expressed autism risk factor transcripts across 21 cell clusters (Wilcox test, adjust p < 0.05, log (fold change) > 0.3) for differentially expressed monogenic autism risk factor transcripts (top panel) and differentially expressed *16p11.2* transcripts (bottom panel).

**Figure 2:**
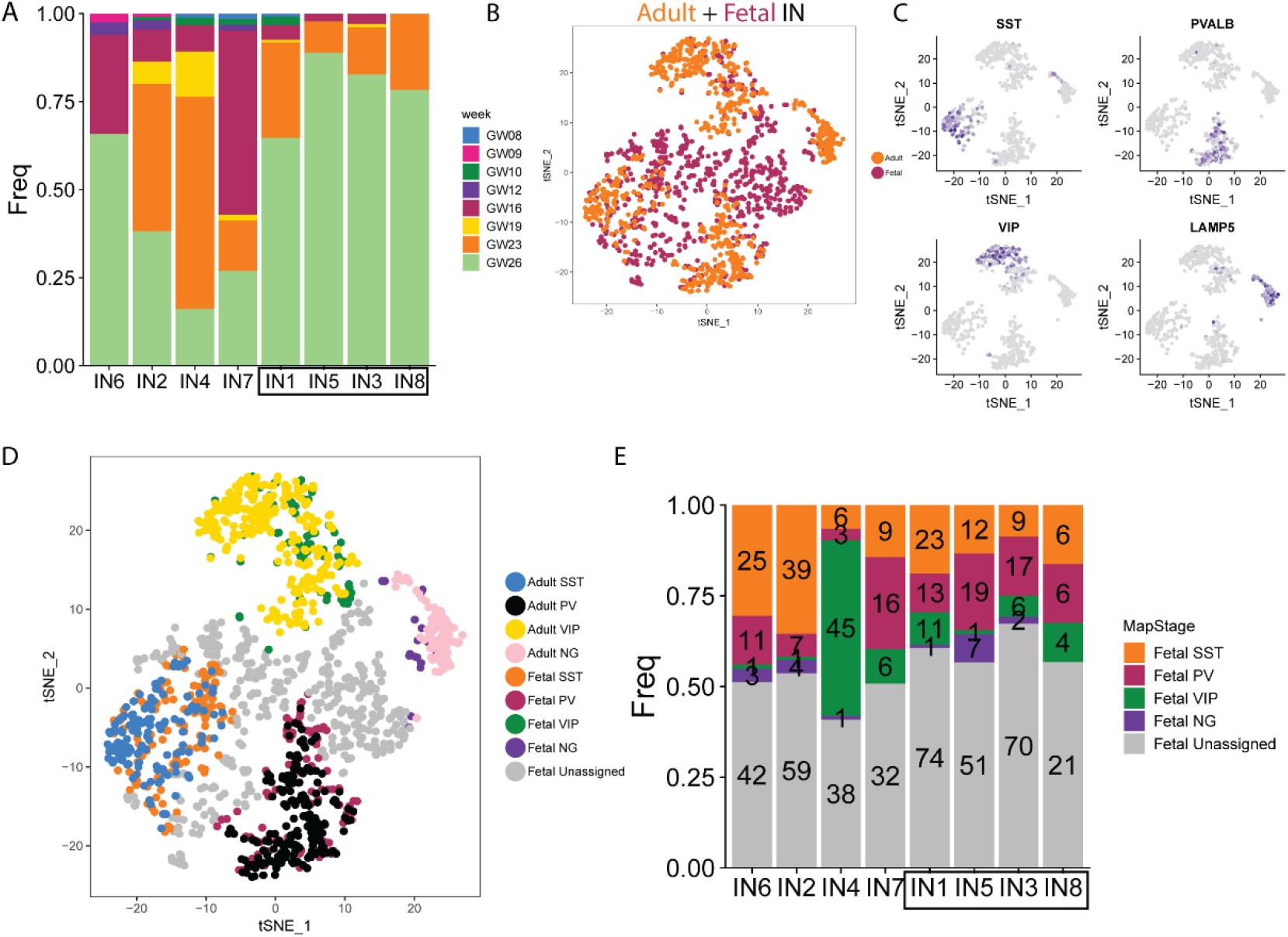
**(A)** Bar plot depicting the proportion of sample ages in each IN cluster. **(B)** Canonical correspondence analysis (CCA) integrating the foetal and adult human scRNA-seq data. **(C)** CCA**-**KNN analysis in t-SNE space provides a method to categorise immature INs into SST, PV, VIP and Neuroglia form (NG) classes according to their transcriptomic similarity to mature neurons from human adult cortex. **(D)** Gradient plots showing gene expression pattern of marker genes of IN lineages in t-SNE space. **(E)** Bar plot depicting the number and proportion of IN cell types in each IN cluster.

For the mouse scRNA-seq datasets at of *Dlx6a-cre* fate-mapped cortical inhibitory neurons, the pre-processed Seurat objects were downloaded from the author’s share link (https://www.dropbox.com/s/qe2carqnf9eu4sd/Filtered_Mayer-et-al.Rda.zip?dl=0) (Mayer et al., 2018). We used the authors’ original classification of seven IN cell types (Sst, Pvalb, Vip, Id2, Nos1, Th and Igfbp6).

All three datasets were converted into Seurat objects by R package Seurat (version 2.3.0) for further analysis. In detail, in the dataset from Zhong et al., raw read counts were normalized based on the original paper described. Any cells with less than 1000 genes expressed were removed, and any gene expressed by less than 3 cells at less than 1 normalized expression value was removed. Pseudogenes, miRNA, rRNA, mitochondrial associated and ribosome related genes were excluded from further analysis. The filtered gene expression matrix and the classification of the cardinal cell classes were used to create a Seurat object. We also create a Seurat object for the dataset from Lake et al. using the same procedure. The pre-processed Seurat object from Mayer et al was not changed. The scRNA-Seq data was also analyzed with BBrowser version 2.2.44 (SingleCell).

### Lists of autism risk genes

Monogenic autism associated genes were downloaded from the SFARI database (released May 2019) (https://gene.sfari.org/database/human-gene/) and the 83 highest ranking (SFARI 1+2) were analysed as these genes are significant statistically in genome-wide studies between cases and controls. Besides these monogenetic genes, the copy number variance (CNV) of genetic loci (CNV genes), either deletions or duplications, are also linked to autism. We selected the 27 protein coding genes at the *16p11.2* locus since both duplication and deletion of these genes has been linked to significantly increased incidence of autism representing a potentially polygenic cause of autism.

### Clustering and visualization of cell types

The identification of six cardinal cell classes were obtained from the original paper and re-plotted in a two-dimensional space of t-Distributed Stochastic Neighbor Embedding (tSNE). In details, the highly variable genes (HVGs) were identified using Seurat function FindVariableGenes. The mean of logged expression values was plotted against variance to mean expression level ratio (VMR) for each gene. Genes with log transformed mean expression level between 1 and 8, and VMR lower than 1.2 were considered as highly variable genes. Then principal component analysis (PCA) was performed with RunPCA function in Seurat using HVGs to analyze all the cells. Following the PCA, we conducted JACKSTRAW analysis with 100 iterations to identify statistically significant (p value < 0.01) principal components (PCs) that were driving systematic variation. We used tSNE to present data in two-dimensional coordinates, generated by RunTSNE function in Seurat, and the first 7 significant PCs identified by JACKSTRAW analysis were used as input to RunTSNE function. Perplexity was set to 20. t-SNE plot and the violin plot were generated using R package ggplot2.

We further clustered the three cardinal cell classes (NPC, ExN and IN) from the foetal cortical dataset. Due to the different number of cells and the variant gene expression pattern in each cardinal cell class, the HVGs were identified using the same method but with the different parameters. For the cells in NPC, genes with log transformed mean expression level between 0.5 and 8, and VMR lower than 1.2 were considered as highly variable genes. For the cells in ExN and IN classes, genes with log transformed mean expression level between 1 and 10, and VMR lower than 0.5 were considered as highly variable genes. Then the statistically significant PCs were calculated by JACKSTRAW analysis and used as input to get tSNE coordinates. Clustering was done with Luvain Jaccard algorithm using t-SNE coordinates by FindClusters function from Seurat. The resolution parameters used to IDENTIFY clusters within the three cardinal cell classes were: NPC, resolution = 1; ExN, resolution = 0.1; and IN, resolution = 0.5. Other parameters that we left at default.

### Identification of differential expressed genes

All differential expression (DE) analyses were conducted using Seurat function *FindAllMarkers*. In brief, we took one group of cells and compared it with the rest of the cells, using Wilcoxon rank sum test. For any given comparison we only considered genes that were expressed by at least 33% of cells in either population. Genes that exhibit p values under 0.05, as well as log fold change over 0.33 were considered significant. All heatmaps of DE analysis were plotted using R package pheatmap (Figure 1C, Figure 3A and 3B, and Figure 5D).

**Figure 3.**
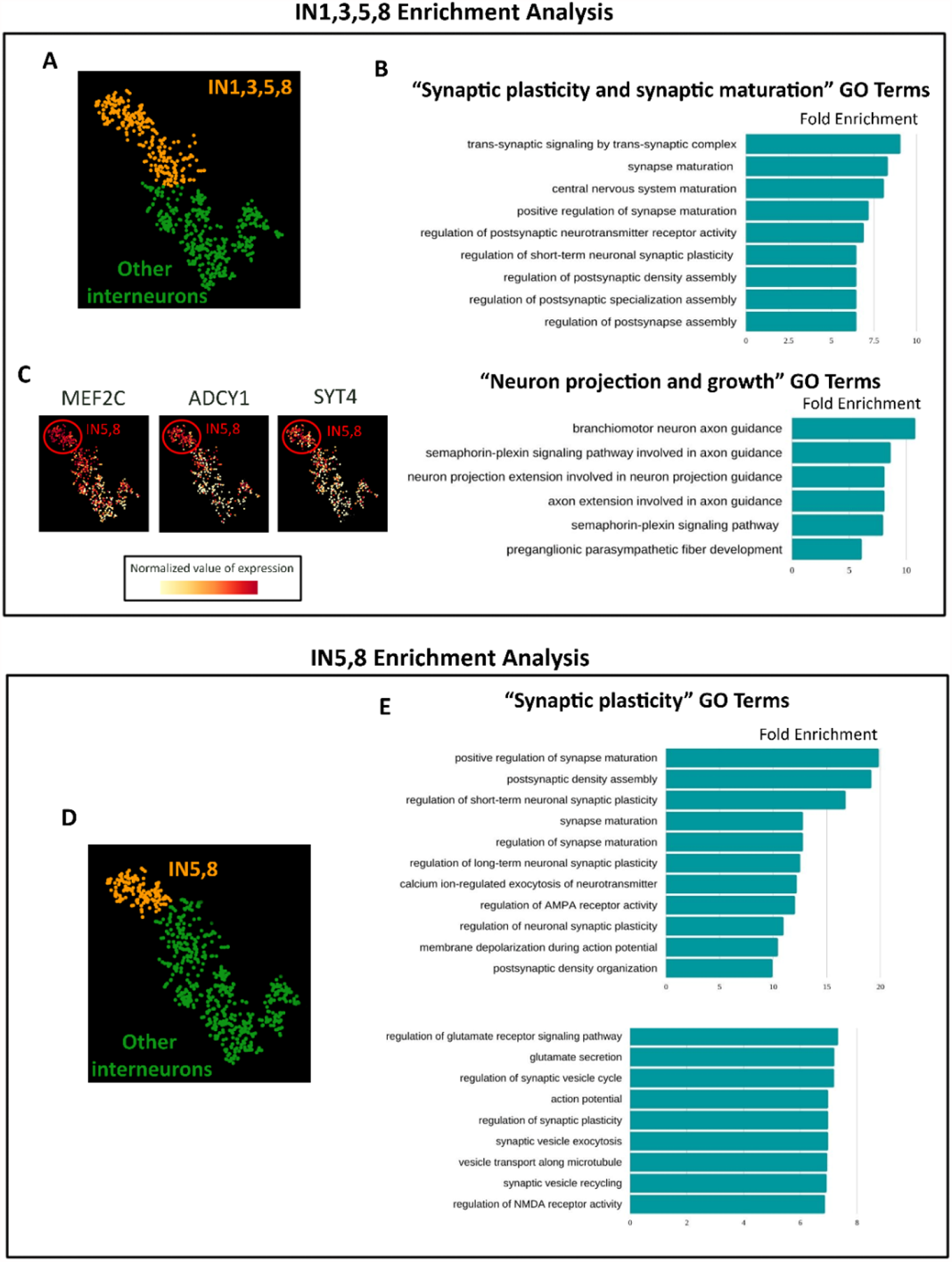
Characterisation of INs by gene ontology analysis (A-C) Gene ontology (GO) analysis in IN1,3,5,8 (orange in A) versus other INs (green in A) reveals enrichment of GO terms associated with (B) synaptic maturation and plasticity and axon extension and guidance. (C) gradient plots of MEF2C, ADCY1, and SYT4 showing that these transcripts are expressed in a gradient across INs with highest expression in IN5,8. (D,E) Gene ontology (GO) analysis in IN5,8 (orange in D) versus other INs (green in D) reveals enrichment of GO terms associated with (E) synaptic maturation and plasticity.

**Figure 4:**
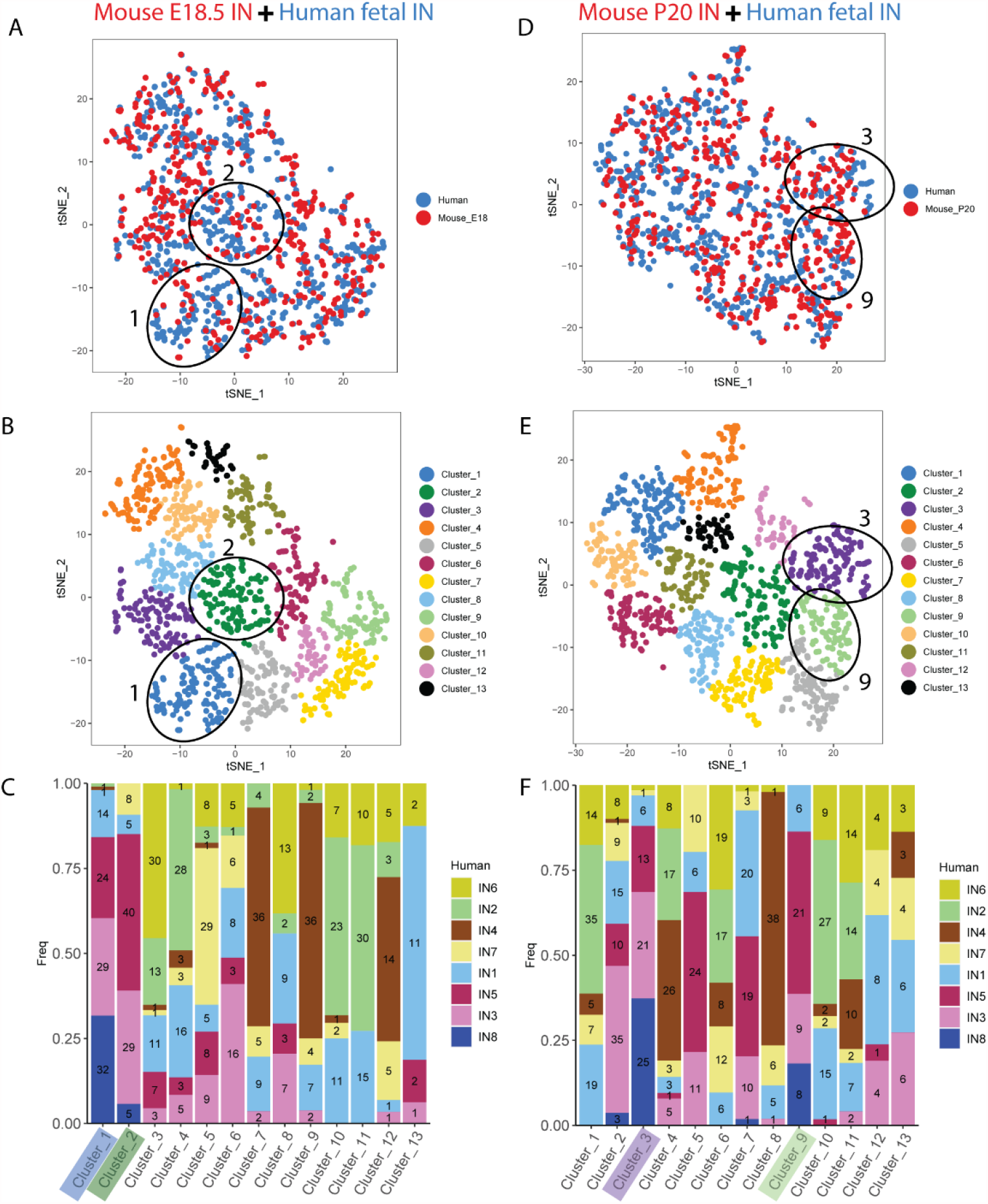
Identifying transcriptomic correlates between developing human and mouse INs at (A-C) E18.5 and (D-F) P20. (A,D) CCA integration of mouse (red) and human (blue) INs in tSNE space. (B,E) JACCARD clustering into 13 mixed clusters. (C,F) Distribution of human IN1-8 INs in each of the mixed clusters with numbers of cells shown within each bar. The mixed clusters 1&2 for E18.5 (A-C) and 3&9 for P20 (D-F) that are most enriched for human IN1,3,5,8 cells are indicated on each panel.

**Figure 5.**
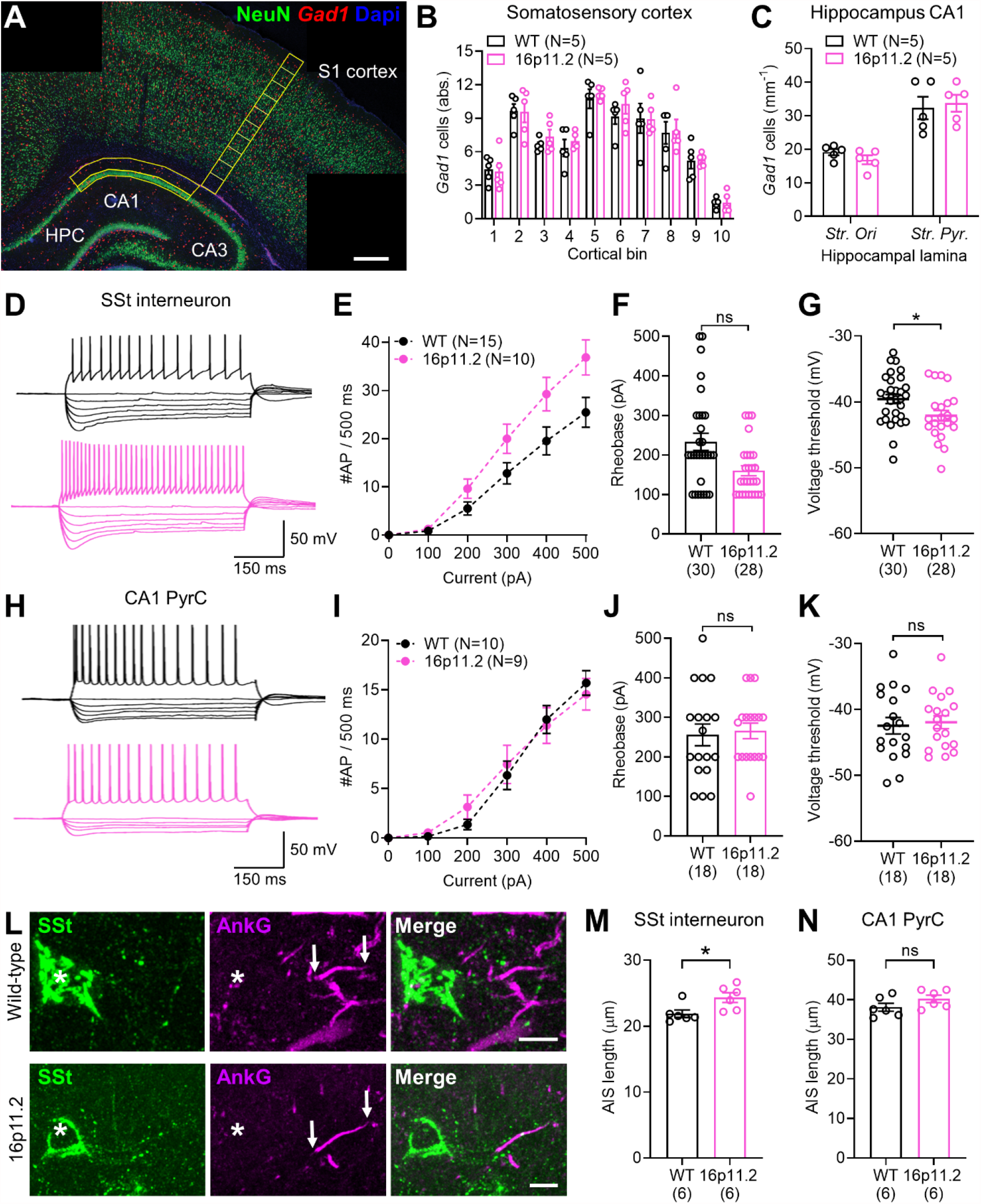
Selective hyperexcitability of SSt INs in *16p11.2*^*+/-*^ rat hippocampus, but with no change in IN number. **(A)** Overview micrograph of *Gad1* mRNA and NeuN protein expression in the rat hippocampus and cortex at P21. Gad1 expressing Ins (red) and NeuN expressing, Gad1 negative excitatory neurons (green) can be observed in the cortex and hippocampus. Scale bars 400µm **(B)** Quantification of *Gad1*-positive IN (IN) number through the somatosensory cortex in WT (black, n=5) and 16p11.2^+/-^ (pink, n=5) rats. Counting areas indicated yellow in A with cortical bins numbered from 1 at the ventricular edge to 10 at the pial surface. **(C)** Quantification of the combined total number of *Gad1*-positive neurons in the s*tr.oriens* (*Str. ori*) and s*tr. pyramidale* (*Str pyr*.) of the CA1 region of the hippocampus in WT (N=5) and 16p11.2^+/-^ (N=5)rats. Counting area indicated in **A**. (**D**) Representative action potential discharge in response to hyper- to depolarising current steps in putative SSt-INs, from the *str. oriens* of CA1 from WT (top) and 16p11.2^+/-^ rats (bottom). **(E)** Summary current-frequency plots from identified SSt INs from WT (N=15 rats) and *16p11.2*^+/-^ (N=10 rats). (**F**) Quantification of rheobase current in identified SSt-INs from WT (n=30 cells) and *16p11.2*^+/-^ (n=28 cells) rats. (**G**) Measurement of the voltage threshold of the first action potential elicited at rheobase for the same cells in **F**. (**H-K**) the same analysis performed in CA1 pyramidal cells from WT (N=10 rats, n=18 cells) and *16p11.2*^+/-^ (N=9 rats, n=18 cells). (**L**) Representative micrographs showing immunohistochemical labelling for SSt (green), the AIS marker AnkyrinG (AnkG, magenta), and their overlap (merge). The SSt soma is indicated with an asterisk (*) and the start and end of the AIS localised to that IN indicated (arrows). Scale bar: 10 µm. (**M**) Quantification of the AIS length of SSt INs from WT (N=6 rats, n= 162 AIS) and *16p11.2*^+/-^ (N=6 rats, n=151 AIS). (**N**) Quantification of AIS lengths of putative CA1 pyramidal cells from WT (N=6 rats, n= 150 AIS) and *16p11.2*^+/-^ (N=6 rats, n=155 AIS) rats. Statistics shown: ns – p>0.05, * - p<0.05, from Linear Mixed Effects modelling.

### MetaNeighbor analysis

MetaNeighbor analysis was performed using the R function MetaNeighbor with default settings (Crow et al., 2018). The AUROC (Area under the Receiver Operating Characteristic) scores produced by MetaNeighbor analysis indicate the degree of correlation between cell clusters. Three gene lists were used as input to do MetaNeighbor analysis among the 21 clusters of human foetal dataset: Highly variable genes (HVGs) identified as significant differentialy expressed genes (DEGs) between the clusters (Figure S2A); monogenic autism risk genes (Figure S2B); and*16p11.2* genes (Figure S2C). The results from the MetaNeighbor analysis were plotted as a heatmap using the gplots function heatmap. For a given gene set each pairwise comparison between cell clusters is given an AUROC score ranging from 1.0 (red on the heatmap) indicating that cells were highly probable to be of the same type to 0.0 (blue on the heatmap) indicating that it was highly improbable that the cells were of the same type. A score on 0.5 (yellow on the heatmap) indicates that the gene set used was unable to distinguish between the cells better than by chance.

### Projection based on multiple datasets

We conducted canonical correlation analysis (CCA) and *k*-nearest neighbors analysis (KNN) as we previous described to classify the cell types of foetal INs based on the cell type features in the adult transcriptomics datasets (Mi et al., 2018). Briefly, we first performed random forest analysis within HVGs to do feature selection for both foetal and adult human cortical INs. Then we selected the shared HVGs between two datasets that best represented the feature of IN cell types. The HVGs were used as input gene list to RunCCA function, and the first 4 dimensions were used as input to AlignSubspace function. The aligned projection vectors were used as input to do dimensional reduction by RunTSNE function. Perplexity was set to 40. We used the two t-SNE coordinates for adult cells to conduct KNN and re-assign foetal IN identities using the knn.cv function from R package FNN. A foetal IN was assigned the identity represented by the majority, and at least 5, of its closest 30 neighbours; in case of ties, the cell remains unassigned. t-SNE plots, and the bar plots were generated using R package ggplot2.

### Gene ontology analysis

The resulting gene list, ordered by sign-adjusted P value, was the input for gene set enrichment analysis to test for enriched gene ontology (GO) terms using the clusterProfiler package version 3.4.4 with default settings. GO term analysis was performed on three categories (Biological process. Molecular function. Cellular component), and gene sets with a BH adjusted P < 0.05 were considered to be significantly enriched. The top three significant GO terms in each category were plotted by R package ggplot2.

### Animals

All rats were bred in-house according to Home Office UK legislation and licenses approved by the University of Edinburgh Ethical Review Committees and Home Office. Animal husbandry was in accordance with UK Animals (Scientific Procedures) Act of 1986 regulations. Rat 16p11.2 DEL rat model (*16p11.2*^*+/-*^ ) was generated by CRISPR/Cas9 genome editing of the Sprague Dawley line (Qiu et al., 2019). Rats were maintained on the Sprague Dawley background. P21 rat tissue was fixed by transcardial perfusion with 4% paraformaldehyde in PBS, brains were then dissected and immersed in 4% paraformaldehyde in PBS overnight at 4°C.

### In Situ Hybridisation and Immunofluorescence labelling

Brains were cryoprotected in 30% sucrose in PBS, embedded in OCT and sectioned at a thickness of 10µm using a cryostat (Leica, CM3050 S). Frozen sections were then mounted on SuperFrost Plus*™* slides (Thermo Fisher). *Gad1* In situ hybridisation on frozen sections was performed as previously described (Wallace and Raff, 1999). NeuN Immunofluorescence was performed following in situ hybridisation as described previously (Clegg et al., 2014) with rabbit anti-NeuN (1/300, Abcam) Secondary antibodies used were donkey anti-goat Alexa Fluor 488 and donkey anti-rabbit Alexa Fluor 568 (both used 1/200 and from Life Technologies). Tissue was counterstained using DAPI (1/1000, Life Technologies).

Axon initial segment (AIS) labelling, was performed as previously described (Oliveira et al., 2020). Briefly, rats were perfused as described above, then post-fixed for 1 hour at room temperature with 4% paraformaldehyde in 0.1 M phosphate buffer (PB). Brains were then transferred to 0.1 M phosphate buffered saline (PBS) and 60 µm thick coronal slices containing the CA1 region of hippocampus were cut on an oscillating blade vibratome (Leica VT1000, Leica, Germany) and transferred to PBS. Briefly, sections were rinsed in PBS then transferred to a blocking solution containing 10% normal goat serum, 0.3% Triton X-100 and 0.05% NaN_3_ diluted in PBS for 1 hour. Primary antibodies raised against AnkyrinG (1:1000; 75-146, NeuroMab, USA) and somatostatin (Somatostatin-14, T-4102.0400; 1:1000, Peninsula Labs, USA) were applied in PBS containing 5% normal goat serum, 0.3% Triton X-100 and 0.05% NaN_3_ for 24-72 hours at 4 °C. Slices were washed with PBS and then secondary antibodies applied (Goat anti-rabbit AlexaFluor 488 and Goat anti-mouse AlexaFluor 633, Invitrogen, UK, both 1:500) in PBS with 3% normal goat serum, 0.1% Triton X-100 and 0.05% NaN3 added, overnight at 4°C. Sections were then washed with PBS, desalted in PB, and mounted on glass slides with Vectashield^®^ mounting medium (Vector Labs, UK). Confocal image stacks of either the *str. pyramidale* or *str. oriens/*alveus border were acquired on a Zeiss LSM800 laser scanning microscope equipped with a 63x (1.4 NA) objective lens at 1024×1024 resolution (step size of 0.25 µm). Individual AIS were measured offline using ImageJ as segmented lines covering the full extent of AnkyrinG labelling observed. As in SSt INs the AIS often emerges from a proximal dendrite, they were only identified where they emerged from a clearly fluorescent labelled dendrite. A minimum of 25 AIS were measured from each rat.

For identification of somatostatin INs, slices were fixed following whole-cell patch-clamp recording (see below) and fixed overnight in 4% PFA in 0.1 M PB. Immunofluorescent labelling was performed according to the same protocol as above, but excluding the AnkyrinG antibody. Secondary antibodies (goat anti-rabbit AlexaFluor488, 1:500, Invitrogen, Dunfermline, UK) were applied with the added inclusion of fluorescent-conjugated streptavidin (Streptavidin AlexaFluor 633, 1:500, Invitrogen, Dunfermline, UK) to visualise recorded neurons.

### Imaging

All fluorescence images were acquired using either a Leica AF6000 epifluorescence microscope coupled to a Leica DFC360 digital camera running Leica LAS-X software, or a Nikon Ti: E Inverted confocal microscope running Nikon NIS-Elements Confocal software.

### NeuN/Gad1 Cell Quantification

*Gad1*^*+*^ and NeuN^+^ cells within the cortex were quantified by counting *Gad1*^*+*^ (red) and NeuN^+^ cells (green) within a 200µm wide column spanning the somatosensory cortex (indicated region, Figure 5A). *Gad1*^*+*^ and NeuN^+^ cell position was quantified by counting cells in 10 adjacent counting bins within the same 200µm wide column spanning the somatosensory cortex.

*Gad1*^*+*^ and NeuN^+^ cells within the hippocampus were quantified by counting cells within the *str. oriens* and *str. pyramidale* of the CA1 region (indicated region, Figure 5A). To control for the varying size of the counting area *Gad1*^*+*^ cell number was expressed as *Gad1*^*+*^ cells per length (mm) of the CA1 region, length was measured along the centre of the *str. pyramidale*. Gad1^+^ cells were classified as belonging to *str. pyramidale* if in contact with NeuN^+^ ;*Gad1*^*-*^ pyramidal cells, all other hippocampal *Gad1*^*+*^ cells superficial to this layer were classified as belonging to *str. oriens*. All measurements and quantification was performed using FIJI software.

### *In vitro* slice electrophysiology

Acute rat brain slices were prepared as previously described (Oliveira et al., 2021). Briefly, rats were decapitated without anaesthesia and the brain rapidly dissected into ice-cold sucrose-modified artificial cerebrospinal fluid (ACSF; in mM: 87 NaCl, 2.5 KCl, 25 NaHCO_3_, 1.25 NaH_2_PO_4_, 25 glucose, 75 sucrose, 7 MgCl_2_, 0.5 CaCl_2_), which was saturated with carbogen (95 % O2/5 % CO2). 400 μm horizontal brain slices were cut on a vibratome (VT1200S, Leica, Germany) and transferred to sucrose-ACSF at 35°C for 30 min and then room temperature until needed.

For whole-cell patch-clamp recordings slices were transferred to a submerged recording chamber flowing with pre-warmed ACSF (in mM: 125 NaCl, 2.5 KCl, 25 NaHCO_3_, 1.25 NaH_2_PO_4_, 25 glucose, 1 MgCl_2_, 2 CaCl_2_), bubbled with carbogen, and perfused a rate of 4-6 mL.min^-1^ at 30± 1 °C). Slices were viewed under infrared differential inference contrast microscopy with a digital camera (SciCamPro, Scientifica, UK) mounted on an upright microscope (SliceScope, Scientifica, UK) with 40x water-immersion objective lens (1.0 N.A., Olympus, Japan). Recording pipettes were pulled from borosilicate glass capillaries (1.7 mm outer/1mm inner diameter, Harvard Apparatus, UK) on a horizontal electrode puller (P-97, Sutter Instruments, CA, USA), which when filled with a K-gluconate based internal solution (in mM 142 K-gluconate, 4 KCl, 0.5 EGTA, 10 HEPES, 2 MgCl_2_, 2 Na_2_ATP, 0.3 Na_2_GTP, 1 Na_2_Phosphocreatine, 2.7 Biocytin, pH=7.4, 290-310 mOsm) which resulted in a 3-5 M*Ω* tip resistance. Cells were rejected if: they were more depolarised than -50 mV, had series resistance >30 MΩ, or the series resistance changed by more than 20% during the recording. Recordings were performed with a MultiClamp 700B (Molecular Devices, CA, USA) amplifier and filtered online at 10 kHz with the built-in 4-pole Bessel filter and digitized at 20 kHz (Digidata1550B, Molecular Devices, CA, USA).)

Cells were identified either as CA1 pyramidal cells (CA1 PCs) with having large, ovoid somata located in *str. pyramidale* and an apical dendrite entering *str. radiatum* or somatostatin INs having bipolar, horizontally oriented somata at the *str. oriens*/alveus border. All intrinsic membrane properties were measured in current-clamp. Passive membrane properties, included resting membrane potential, membrane time constant, and input resistance, were measured from hyperpolarising steps (−10 pA, 500 ms duration), from resting membrane potential. Active properties were determined from a series of hyper- to depolarising current steps (−500 to +500 pA, 500 ms) from a holding potential of -70mV, maintained with a bias current injection. All AP properties were determined from the first AP elicited above rheobase. Spontaneous EPSCs were measured in voltage-clamp from a holding potential of -70mV and detected offline based on a triexponential curve fit and a threshold of 3*SD of the baseline noise. Traces were collected in pCLAMP 9 (Molecular Devices, CA, USA) and stored on a desktop computer. Analysis of electrophysiological data was performed offline Stimfit (Guzman, Schlögl, and Schmidt-Hieber 2014), blind to both genotype. All data from somatostatin INs is shown only for those cells where clear immunofluorescent labelling was detected at the level of the soma.

### Statistics

All rat experiments and analyses were performed blind to genotype, which were sampled in a random manner between experimental days. All data shown as mean ± standard error of the mean (SEM), with the number of cells (n) and animals (N) indicated where appropriate. All electrophysiology data are reported as cell averages. All histology data (AIS lengths and cell counts) are shown as animal averages. Minimum sample size was calculated based on our previous effect size for cellular hyperexcitability and AIS length (Booker *et al*., 2020), assuming 80% power to determine 95% probability of rejecting the null-hypothesis. Statistical comparisons were performed using a linear mixed-effect model (or its generalised form) using the *lme4* package in R (Bates *et al*., 2015), with genotype or cell-type as fixed effect, with slice/animal/litter included as random effects. Based on the linear mixed-effects model, p-values for statistical effects were tested using the Wald test, based on effect size and variance determined from the relevant mixed-effects model. For experiments examining the density of interneurons and principal cells, animal average densities were the principal replicate which was tested with 2-way ANOVA. Statistical significance was assumed if p<0.05.

## RESULTS

### Differential expression of autism risk genes among foetal cortical cardinal cell classes

A general supposition is that functional disruption of a gene will more likely affect the cells expressing high levels of it’s transcript. Based on such a principle, a cell type expressing high levels of an autism associated transcript is regarded as potentially vulnerable to genetic mutation in that gene with the resulting cellular phenotype contributing to the development of autism. Accordingly, we have calculated differential expression of autism associated transcripts among cell types in foetal human cerebral cortex to identify cells potentially vulnerable to autism genetic risk factors during brain development.

We started with a scRNA-seq dataset comprising 2306 cells taken from human foetal pre-frontal cortex spanning gestational weeks (GW) 8 to 26 (Zhong et al., 2018). Six cardinal cell types were identified in the authors’ original classification, including neural progenitor cells (NPCs), excitatory neurons (ExN), interneurons (IN), oligodendrocyte precursor cells (OPCs), astrocytes, and microglia (Fig. 1A,B and Fig. S1A,B). Differentially expressed genes (DEGs) were calculated across these six cardinal cell classes. Based on the DEGs, we find that the six cell classes showed distinct cardinal class aggregation and specific gene expression profiles associated with neural progenitor cells (NPC), excitatory neurons (ExN), inhibitory neurons (IN), oligodendrocyte precursors (OPC), astrocytes, and microglia. A list of well-known cell class markers that are included in the DEGs was used to illustrate the classification across six cardinal cell classes (Nowakowski et al., 2017, Pollen et al., 2014, Camp et al., 2015) (Fig. 1B). The markers used to identify different cardinal cell classes were: *PAX6, HES2* and *VIM* (NPCs); *NEUROD2, NEUROD6* and *RBFOX1* (ExNs); *GAD1, GAD2, DLX1* and *DLX2* (INs); *OLIG1, OLIG2* and *COL20A1* (OPCs); *GFAP, AQP4* and *SLCO1C1* (astrocytes); *PTPRC* and *P2RY12* (microglia). The expression pattern of these marker genes show that the cardinal cell classes were correctly represented in our analysis (Fig 1B).

Then we identified the expression pattern of autism risk factor transcripts across the cardinal cell classes and found that 17/83 high confidence and strong candidate monogenic risk factor transcripts and 7/27 *16p11.2* transcripts were significantly differentially expressed between cardinal cells classes (Fig 1C). A heatmap of expression of the monogenic autism risk factor transcripts (Fig 1C – top) and the *16p11.2* transcripts (Fig 1C – bottom) shows expression of each autism risk factor transcript (rows) in each of the six cardinal cell classes (columns). Transcript levels with expression greater than average across the cardinal classes are shown in red, while transcripts with lower than average expression are shown in blue. There was no obvious pattern to suggest that any cardinal class was particularly vulnerable to a large proportion of either monogenic autism genetic risk factors.or the *16p11.2* microdeletion.

### Identification of human foetal INs potentially disproportionately vulnerable to genetic autism risk factors

The single-cell approach allows us to investigate the variability of highly expressed genes among molecularly defined cell subpopulations and identify cells within cardinal classes which may be vulnerable to genetic autism risk factors. Based on unsupervised clustering, we subdivided the cardinal classes into 21 different cell clusters (Fig. 1D): 6 for NPCs (P1-6); 4 for ExNs (N1-4); and 8 for INs (IN1-8). The non-neuronal cardinal cell classes (OPC, astrocytes, and microglia) contained small numbers of tightly clustered cells and were not further subdivided.

We clustered the 21 clusters according to transcriptomic similarity using MetaNeighbour analysis with ∼2000 highly variable genes (Fig. S2A) and used this ordering to generate a heatmap of the 62/83 monogenic risk factor transcripts (Fig. 1E, top) and 17/27 *16p11.2* transcripts genes (Fig. 1E, bottom) that were significantly differentially expressed between clusters. Violin plots of all autism risk transcripts (including those not differentially expressed between clusters) for the 83 monogenic autism risk transcripts (Fig. S3A) and 27 *16p11.2* transcripts (Fig. S3B) show the expression profile in each cluster.

Of the differentially expressed monogenic risk factor transcripts (Fig 1E, top) a few were enriched in progenitor cells, (for example *IRF26PL, BCL11A*, and *CHD2*), with fewer transcripts enriched in excitatory neurons (for example *TBR1*). Other transcripts were expressed across many clusters (for example *KMT2A, ILF2, SMARCC2, SRSF11, UPF3B, TNRC6B*), while others showed relatively low expression across cell types (for example *GRIN2B, MAGEL2, and MET*). A striking feature of the heatmap was a preponderance of relatively high gene expression (red shading) in the IN cell-types (for example *SCN2A, SCN9A, DEAF1, SHANK2, RIMS1, GRIP1, SYNGAP1*), which was most apparent for the IN subgroups IN1,3,5,8 (purple highlight in Fig. 2E and circle in Fig. 1D) compared to IN2,4,6 7 (green highlight in Fig. 2E and circle in Fig. 1D) and other clusters.

We observed a similar pattern for *16p11.2* transcript expression (Fig 1E, bottom) with a preponderance of high expression in IN clusters1,3,5,8, indicating specifically enriched expression of subset of *16p11.2* transcripts (for example *SEZL6L2, PRRT2, QPRT*, and *YPEL3*) with the next highest number of enriched transcripts in progenitor clusters (for example *KIFF22* and *PPC4C*). Other transcripts are expressed both in progenitors and INs but much less in excitatory neurons (for example *MAPK3*), and others broadly across all cell clusters (for example *TMEM219*) or at very low levels in any cluster (*ASPHD1, C16orf54, C16orf92, SPN, TBX6, ZG16*).

To investigate how well the expression of autism risk factor transcripts defined the cell clusters more systematically we used MetaNeighbor analysis which reports on how similar cells are to each other based on expression of a given input gene set using AUROC scores (Crow et al., 2018). From this, we generated a pairwise comparison matrix between the 21 cell clusters (Fig. S2). Performing MetaNeighbor with the same ∼2000 DEGs used to perform hierarchical clustering (Fig. S2A) we confirmed that cells in each cardinal class were generally more similar within class (red on heatmap) and less similar (blue on heatmap) between cardinal classes. Nevertheless, some neurons (N1 and N2) were quite similar to progenitors (P1-P6) likely indicating that they represented relatively immature excitatory neurons that retained some progenitor identity. Within the INs there was a clear divide between IN1,3,5,8 (red box in Fig. S2A-C) and IN2,4,6 (green box in Fig. S2A-C) cluster groups, with highest similarity within group and low similarity between groups. Next, we performed the same analysis for the 83 SFARI monogenic autism risk factor transcripts (Fig. S2B) and found that strongest similarity was retained within the progenitor group P1-P6 and the IN group IN1,3,5,8. A similar pattern emerged when we used the 27 *16p11.2* transcripts as the gene-set (Fig. S2C), although here the strongest similarity was within the progenitor group P1-6 and between IN5 and IN8. These results indicate that the immature INs can be robustly distinguished from other cells in the developing brain by their expression pattern of autism risk factor transcripts.

We conclude from the gene expression analysis that a subset of developing INs, IN1,3,5,8, in GW8-26 human foetal cerebral cortex express a high proportion of autism associated transcripts at higher levels than other cells. This suggests that IN1,3,5,8 are vulnerable, and in some instances selectively vulnerable, to a large number of independent monogenic genetic autism risk factor and *16p11.2* microdeletion during cerebral cortex development.

### Properties of human foetal INs potentially vulnerable to autism risk factors

Having identified IN1,5,3,8 as a potentially important class of INs targeted by genetic autism risk factors we next examined their developmental and transcriptional properties. We found that very few IN1,3,5,8 INs were present during the earlier stages (GW8-19) of cerebral cortex development, typically appearing from GW23 and with the vast majority of IN1,5,3,8 INs present at GW26 (Fig. 2A). On the other hand, IN2,4,6,7 clusters contained higher proportions of cells from earlier stages (Fig. 2A), suggesting that IN1,3,5,8 might represent a more mature state than the rest of IN clusters in our dataset. INs have reached the cortex in substantial numbers by GW16 (Fig. S1B) indicating that IN1,3,5,8 cells correspond to a stage of the developmental trajectory after tangentially migrating INs enter the cortex (Hansen et al., 2013; Ma et al., 2013).

We used canonical correlation analysis (CCA-KNN) to integrate foetal (Zhong et al., 2018) and mature (Lake et al., 2018) human IN scRNAseq datasets to identify mature cell types corresponding to developing IN1,3,5,8 clusters. We first reduced the dimensionality of both datasets (adult IN cells = orange; foetal IN cells = blue) onto the same two-dimensional space using *t*-SNE (Fig. 2B), which allowed the identification of 4 major cell types of adult INs based on the expression of variable genes shared between both datasets. We then assigned identities to adult based on expression of markers for PV, SST, VIP and neurogliaform cells illustrated by gradient plots of *SST, PVALB, VIP* and/or *LAMP5* transcripts to identify defined classes of cortical INs (Fig. 2C). This then allowed us to assign foetal INs to each of these cell types based on transcriptional similarity to the mature INs (Fig. 2D). Of the IN1,3,5,8 cells classified in this manner we found that they were not homogenous, but rather consisted of PV, SST, and VIP cell types (Fig. 2E). A parsimonious interpretation is that the foetal IN1,5,3,8 are cells destined to become several categories of mature IN cell types, although this awaits further investigation as assignation of cell lineage from scRNA-seq data is ambiguous.

To gain further insight into the identity and developmental cell state represented by IN1,3,5,8, we first performed differential expression analysis with respect to other INs (Fig. 3A) and used the genes enriched in IN1,3,5,8 with a log fold change higher than 3 (1623 genes FDR < 0.001) to test for Gene Ontology (GO, biological process) enrichment. We found that within the top 30 GO terms (ordered according to Fold Enrichment), 9 categories were related to synaptic plasticity, synaptic maturation and synaptic transmission and 6 categories were related to neuron projection and growth (Fig. 3B), with fold enrichments ranging from 6 to 10, suggesting that IN1,3,5,8 cells show earlier maturation of neurites and synapses than other INs. The Gene Ontology term “Regulation of Synaptic Plasticity” contains 192 genes, from which 54 are differentially expressed in IN1,3,5,8. A closer inspection of the expression pattern in the t-SNE space showed that many of these genes followed a general expression level gradient trend with its maximum levels in INs corresponding to IN5 and IN8 (*MEF2C, ADCY1*, and *SYT4* shown as examples in Fig. 3C). This suggested that the INs in the dataset must be ordered in the t-SNE space according to a gradient of synapse formation, with IN5,8 being the higher extreme of this axis. To confirm this, we performed differential expression analysis between IN5,8 IN cells versus all other INs (Fig. 3D) and found a high enrichment of synaptic plasticity-related terms but this time showing fold enrichments ranging from 7 to 20 (Fig. 3E), almost doubling the values of the previous comparison.

Finally, to gain further insight into IN5,8 neuronal identity, we compared IN5,8 cluster versus all other neurons (including both excitatory and inhibitory, Fig. S4A). Interestingly, enriched functional terms were mainly related to synaptic plasticity, learning and social behaviour (Fig. S4B). Visual inspection of gradient plots in the t-SNE space confirmed that many of the genes linked to synaptic plasticity and maturation are selectively expressed in IN5,8 (Fig. S4C). Together this analysis suggests that IN1,3,5,8 are relatively differentiated INs elaborating processes and forming synapses.

This raises the possibility that by targeting the stage of the IN developmental trajectory represented by IN1,3,5,8, multiple genetic autism risk factors perturb the development of the physiological properties of foetal INs. The remaining interneurons IN2,4,6,7 appear less vulnerable and may represent an earlier stage in the IN developmental trajectory or correspond to different IN lineages. Either way our analysis suggests genetic autism risk factors impact on many INs during foetal development and may affect their function into postnatal life.

### Conservation of potentially vulnerable INs between humans and rodents

These analyses of human foetal neurons suggest the testable hypothesis that a large proportion of autism related genes selectively regulate IN development in the human foetal cortex. A prediction of this hypothesis is that there will be IN phenotypes initiated during human brain development *in utero* that persist into postnatal life and predispose to autism and its comorbid conditions. Such investigation is currently not possible, however, rodent models provide a complementary means to test cellular vulnerability to autism genetic risk factors under physiological conditions. As such, we next confirmed that developing rodent brain contains INs with similar molecular properties to the potentially vulnerable human foetal INs IN1,3,5,8 identified above.

We identified two mouse scRNAseq data sets comprising FACS sorted cortical INs at embryonic day (E) 18.5 and postnatal (P) day 20, when INs are differentiating and forming circuits (Mayer et al., 2018). For each mouse developmental stage we used CCA-KNN to integrate the mouse and human INs into the same tSNE space (Fig. 4 A,D) to allow us to identify mixed clusters of transcriptomically similar mouse and human INs (Fig. 4B,E).

For each mouse developmental stage, we examined how the human IN cell types IN1-8 were distributed between the mixed mouse+human clusters (Fig. 4C,F). This analysis revealed that E18.5 clusters 1 and 2 (indicated in Fig 4A-C) and P20 clusters 3 and 9 (indicated in Fig 4D-F) contained the greatest enrichment of human IN1,3,5,8 cells. Critically, these clusters contained comparable numbers of mouse and human cells indicating that the developing and postnatal mouse possesses INs molecularly similar to human IN1,3,5,8 cells. These findings suggest that INs we have identified as potentially vulnerable to genetic autism risk factors are shared between humans and rodents allowing us to investigate them under physiological conditions using rodent models.

### Changes to IN function in the rat model of *16p11.2* microdeletion

The *16p11.2* microdeletion causes *16p11.2* transcript levels to be reduced by about 50% in humans and rodent models (Tai et al., 2016) (Pucilowska et al., 2015) (Horev et al., 2011) (Blumenthal et al., 2014). As multiple *16p11.2* transcripts are normally highly enriched in developing INs (Fig. 1), we hypothesised that their simultaneous reduced expression following *16p11.2* microdeletion may synergistically impact IN development with post-natal consequences on IN phenotypes. We next set out to test this hypothesis by performing electrophysiological and anatomical interrogation of the rat *16p11.2* microdeletion model (*16p11.2*^+/-^ rats).

As *16p11.2* transcripts are expressed in the ganglionic eminences where IN progenitors reside (Morson et al., 2021), we first asked if the numbers of inhibitory and/or excitatory neurons populating the cortex post-natally was different in the *16p11.2*^*+/-*^ rats. To investigate this, we counted these cardinal cell classes in WT and *16p11.2*^*+/-*^ rats at postnatal day (P) 21, an age by which INs have migrated into the cortex and assumed their final laminar positions forming circuits with resident excitatory neurons. We combined immunostaining for the pan-neuron-specific marker NeuN and *in situ* hybridization for the IN marker *Gad1* to identify NeuN^+^ ;*Gad1*^-^ excitatory neurons and NeuN^+^ ;*Gad1*^+^ INs in the cortex (Fig. 5A). Quantification within a 200µm wide column spanning the somatosensory cortex and hippocampus (shown figure 5A) shows no significant difference between WT and *16p11.2*^*+/-*^ rats in total IN number (*t*_(8)_ = 0.27, *p* = 0.80 *t* test, Figure S5A) or in the proportion of the neuronal population identified as inhibitory or excitatory (*t*_(8)_ = 0.15, *p* = 0.89 *t* test, Fig. S5B). To assess whether the cortical laminar position of INs was altered in *16p11.2*^*+/-*^ rats we quantified IN and excitatory neuron number in 10 adjacent counting areas spanning the somatosensory cortex. We found no significant change in the distribution of INs across the cortex (Figure 5B), nor did we see any change in the inhibitory/excitatory proportion in any counting area (Fig. S5B). Next, we examined IN number and position in the CA1 region of the hippocampus. Total combined IN number within the *str. oriens* (SO) and *st. pyramidale* (SP) of CA1 was unchanged between WT and *16p11.2*^*+/-*^ rats (*t*_(8)_ = -0.09, *p* = 0.93 *t* test, Fig. S5A). Total IN number within the SO and SP was also unchanged indicating that the position of INs within the hippocampus is unaffected in *16p11.2*^*+/-*^ rats (Figure 5C). These data indicate that the *16p11.2* microdeletion does not have a major impact on number or distribution of inhibitory or excitatory neurons in the cerebral cortex or CA1 INs post-natally.

Our bioinformatic analysis indicated that human foetal IN1,3,5,8 cells contribute to mature somatostatin (SST) INs (Fig 2E). As such, we next asked whether the *16p11.2* microdeletion might have an effect on SSt IN development to alter their physiological properties. To test this, we performed whole-cell patch-clamp recordings from identified SSt INs and local pyramidal cells (PCs) in the CA1 region at P21.

We performed recordings from identified INs that expressed SSt in CA1 of the hippocampus, which also show expression of PV in parvalbumin (PV) in 50% of cells (Booker and Vida, 2018). The recorded Sst INs predominantly had horizontally oriented dendrites in *str. oriens*, which when present had an axon extending into *str. lacunosum-moleculare* (Booker and Vida, 2018). In the present study, we recorded from 30 WT (from 15 rats) and 28 *16p11.2*^*+/-*^ (10 rats) SSt INs, which all displayed clear immunoreactivity for the SSt neuropeptide at the somata. In response to depolarisingcurrent injections (0-500 pA, 100 pA steps, Fig. 5D), SSt INs generally responded with high action potential discharge rates, which had a peak action potential discharge of 27.4 ± 2.5 action potentials/500 ms (Fig. 5E). We found that SSt INs from the *16p11.2*^*+/-*^ showed elevated action potential discharge, compared to WT controls (F_(5, 58)_ =5.10,P=0.0006 for interaction of genotype and current, N=15 WT and 10 *16p11.2*^+/-^ rats, Fig. 5E), indicating cellular hyperexcitability. Comparison of the intrinsic physiology of SSt INs revealed a general 17% reduction of rheobase current in 16p11.2^+/-^ compared to WT rats, albeit not significantly so (*p*=0.1087 LME, Fig. 5F), this was accompanied by a 6% hyperpolarisation of the voltage threshold (*p*=0.0279 LME, Fig. 5G). All other physiological parameters were similar between genotypes (Supplementary Table S1).

To confirm that SSt IN intrinsic excitability changes were not a result of compensation mechanism to altered synaptic input from local CA1 PCs (Booker et al., 2020), we asked whether the spontaneous excitatory postsynaptic currents (EPSC) they receive were different between genotypes. We saw no change in either the spontaneous EPSC amplitude (*p*=0.4495, LME) or frequency (*p*=0.2131, LME), implying typical circuit integration of SSt INs to the local network.

We confirmed that the effect on intrinsic cell excitability were restricted to SSt INs, by performing recordings from local ExNs – the CA1 PCs. CA1 PCs were identified on the basis of having a somata located in *str. pyramidale*, with a single large-calibre apical dendrite entering *str. radiatum* as observed under IR-DIC. We obtained recordings from 18 putative CA1 PCs per group from 10 WT and 9 *16p11.2*^*+/-*^ rats. In response to depolarising current injections (0-500 pA, 100 pA steps, Fig. 5H), we observed no change in the number of action potentials generated by the recorded CA1 PCs in the *16p11.2*^*+/-*^ rats compared to WT (F_(5, 85)_=0.9185, P=0.4731, 2-way ANOVA for current/genotype interaction, Fig. 5I). CA1 PCs typically had lower peak action potential discharge rates than for SSt INs (WT: *p*=3.29×10^−5^, *16p11.2*^*+/-*^: *p*=1.63×10^−6^ ; LME). Consistent with this lack of altered action potential discharge, we found no change in CA1 PC rheobase current (*p*=0.7098, LME, Fig. 5J), action potential threshold (*p*=0.4116, LME, Fig. 5K), or any other parameter tested (Supplementary Table S1). These data strongly suggest that excitatory CA1 PCs are physiologically typical in the *16p11.2* rat model of ASD.

We have recently shown that changes to cell excitability effected the voltage threshold and action potential discharge in genetic models of intellectual disability can result from changes to the structure of the axon initial segment (AIS) (Booker et al., 2020). To determine if the changes in SSt IN excitability arise from changes to AIS structure we next performed immunolabelling of perfusion fixed tissue from the hippocampus of WT (N=5) and *16p11.2*^*+/-*^ (N=5) rats. Immunofluorescent double labelling with AnkyrinG reliably labelled the AIS of all neurons in the CA1 region, which could be identified emerging from the soma, or more often the proximal dendrites of immunolabelled SSt INs (Fig. 5L). Comparison of AIS lengths on SSt INs revealed an 11% longer AIS in 16p11.2^+/-^ rats compared to WT (*p*=0.0064, LME, Fig. 5M). There was no change in the lengths of putative CA1 PC AISs between genotypes (*p*=0.0962, LME, Fig. 5N).

Together, consistent with the hypothesis that the *16p11.2* microdeletion selectively targets INs, these data show a preferential increase of SSt IN excitability in the *16p11.2* rat autism model, with no changes observed in the local excitatory principal cells. This increased cellular excitability coincided with selective alteration to the length of the AIS, corresponding to changes in voltage threshold. Together, these changes could potentially lead to an aberrant network activity and gating of information flow through hippocampal circuits.

## DISCUSSION

This study reveals that molecularly defined classes of INs in the foetal human cerebral cortex display enriched expression of multiple gene transcripts associated with autism. This result is striking, suggesting the testable hypothesis that some INs are disproportionately vulnerable to autism genetic risk factors. Within INs as a whole the most autism associated transcripts are enriched in a subset we described as ‘IN1,3,5,8’ suggesting that these cells may represent a convergent cellular target underpinning genetic predisposition of the developing brain to autism and its co-occuring conditions during post-natal life. This poses the question of what is the identity of these cells? Our data-set was acquired from dissected human foetal cortex, which includes INs that are migrating tangentially through and in the cerebral cortex, but not INs undergoing neurogenesis or early migration in the ganglionic eminences (Hansen et al., 2013; Ma et al., 2013). We examined human foetal cortical cells spanning the interval GW8-26 and found IN1,3,5,8 cells are not present in the cerebral cortex before GW23 and then increase in numbers to GW26. As many INs have migrated into the cortex well before GW23 this suggests that IN1,3,5,8 represent a relatively differentiated stage on the IN developmental trajectory. This is consistent with the enrichment of GO terms relating to synapse maturation and neurite formation in these cells. Transcriptomic similarity between foetal IN1,3,5,8 and adult PV, SST, and VIP INs suggests these cells are destined to form a variety of IN cell types and that changes in their developmental trajectory caused by autism causing mutations may have far reaching consequences for the formation on inhibitory circuitry in the post-natal brain.

Although the most striking enrichment of autism associated transcripts was observed in INs we also saw enrichment of some transcripts in other cardinal cell classes. Progenitor cells had a much smaller number of enriched autism associated transcripts than INs and most of these (eg *ADNF, ZNF462, PHIP, HNRNPU, RPS6KA3* for monogenic and *PAGR1, HIRIP3, KIF22*, and *PP4C* for *16p11.2* transcripts). This suggests that progenitor cells in the cerebral cortex may also be vulnerable to a subset of autism causing mutations. Progenitors in the developing neocortex are destined to differentiate into excitatory pyramidal neurons and non-neuronal cell-types (eg astrocytes) suggesting that mutations in these progenitor enriched genes may dysregulate their production or function. This study also prompts future investigation into the expression of autism associated transcripts in IN progenitors located in the ganglionic eminences and the consequence of mutation for IN neurogenesis in humans although our findings in the rat *16p11.2* microdeletion rat model suggest that gross IN output is not affected in this context in rodents.

We found that large numbers of monogenic autism risk factor transcripts are highly expressed in IN1,3,5,8 INs, suggesting that their mutation may contribute to the aetiology of autism via alterations to IN development. This hypothesis remains to be tested for the majority of genes. However, for *ARID1B, DYRK1A, MECP2*, and *CNTNAP2* there is already evidence that monogenic mutation causes abnormal numbers or physiological properties of INs in rodent models (Gao et al., 2018; Jung et al., 2017; Penagarikano et al., 2011; Souchet et al., 2019; Tomassy et al., 2014; Vogt et al., 2018). We also found *KCTD13, MAPK3*, and *MVP* transcripts expressed from the *16p11.2* locus are enriched in IN1,3,5,8. *KCTD13* modulates synaptic transmission by suppressing RHOA signalling via interaction with the ubiquitin ligase *CUL3*, itself an autism risk factor (Escamilla et al., 2017; Willsey et al., 2013). *CUL3* is co-expressed with *KCTD13* in IN1,3,5,8 cells suggesting a molecular mechanism for the *16p11.2* microdeletion to impact on cellular and synaptic function via perturbed RHOA signalling. Interestingly, the inhibition of RHOA pathway has been prosposed as a treatment to restore cognition in *16p11.2* mouse models (Martin Lorenzo et al., 2021). *MAPK3* and *MVP* are both implicated in ERK signalling which impacts diverse cellular processes including cell proliferation, migration, and synaptic function. Indeed, a mouse model of *16p11.2* microdeletion shows elevated ERK signalling leading to perturbed cortical development and autism-like phenotypes (Pucilowska et al., 2018; Pucilowska et al., 2015), although the involvement of INs was not tested.

Our analysis of the *16p11.2* microdeletion rat indicates that neither ExN or IN number and location were altered in the cerebral cortex or hippocampal region CA1, so it seems unlikely that a numerical excitation/inhibition imbalance is present in the *16p11.2*^+/-^ rat model. However, whole-cell patch-clamp recordings in CA1 revealed intrinsic hyperexcitability of SSt INs, coincident with increases AIS length. No effect was observed in CA1 excitatory neurons. This suggests a mechanism by which the *16p11.2* microdeletion perturbs the E/I balance by selectively altering the intrinsic excitability of INs. Although IN hyperexcitability may lead to greater inhibition within cortical circuits and tilt the E/I balance towards inhibition, the complexity of such circuits interaction makes prediction of the consequences to gross activity difficult. However, SSt INs themselves possess both direct inhibitory and disinhibitory mechanisms within the prototypical CA1 circuit, leading to alterations to synaptic plasticity when measured at the circuit level (Leao et al., 2012). As such, the outcome of greater SSt IN activity may directly lead to altered cognition observed in ASC/ID. Furthermore, a shift in the E/I balance has been identified in the somatosensory cortex of ASC mouse models (including *16p11.2*^+/-^ mice), attributed to homeostatic regulation of IN function (Antoine et al., 2019). Our combined bioinformatic and physiological approach diverges from this view, suggesting that SSt INs in *16p11.2* microdeletion are genetically cued to perturbation from early in their developmental trajectory. A deeper understanding of the functional consequences of SSt hyperexcitability on the E/I balance in cortical circuits and their homeostasis requires further investigation.

To conclude, our bioinformatic analysis of developing human foetal cerebral cortex single cell transcriptomes suggests that developing INs are disproportionately vulnerable to genetic autism risk factors, which is supported by physiological correlates in a *16p11.2* microdeletion rat model. This study paves the way for more in depth investigations of how polygenic and monogenic autism risk factors impact on IN development and function.

## Contributions and acknowledgements

YY and IQ performed bioinformatic analysis of human data, SAB, JMC, and AS performed rat experiments and analysis, ZK and OD contributed to linear modelling, SM and YH provided the *16p11.2* microdeletion rats, PCK and DJP contributed to design, TP designed the study and wrote the paper. This work was funded by the Simons Initiative for the Developing Brain (SFARI - 529085) and Biotechnology and Biological Sciences Research Council (BB/M00693X/1).

## SUPPLEMENTARY FIGURES

**Figure S1.**
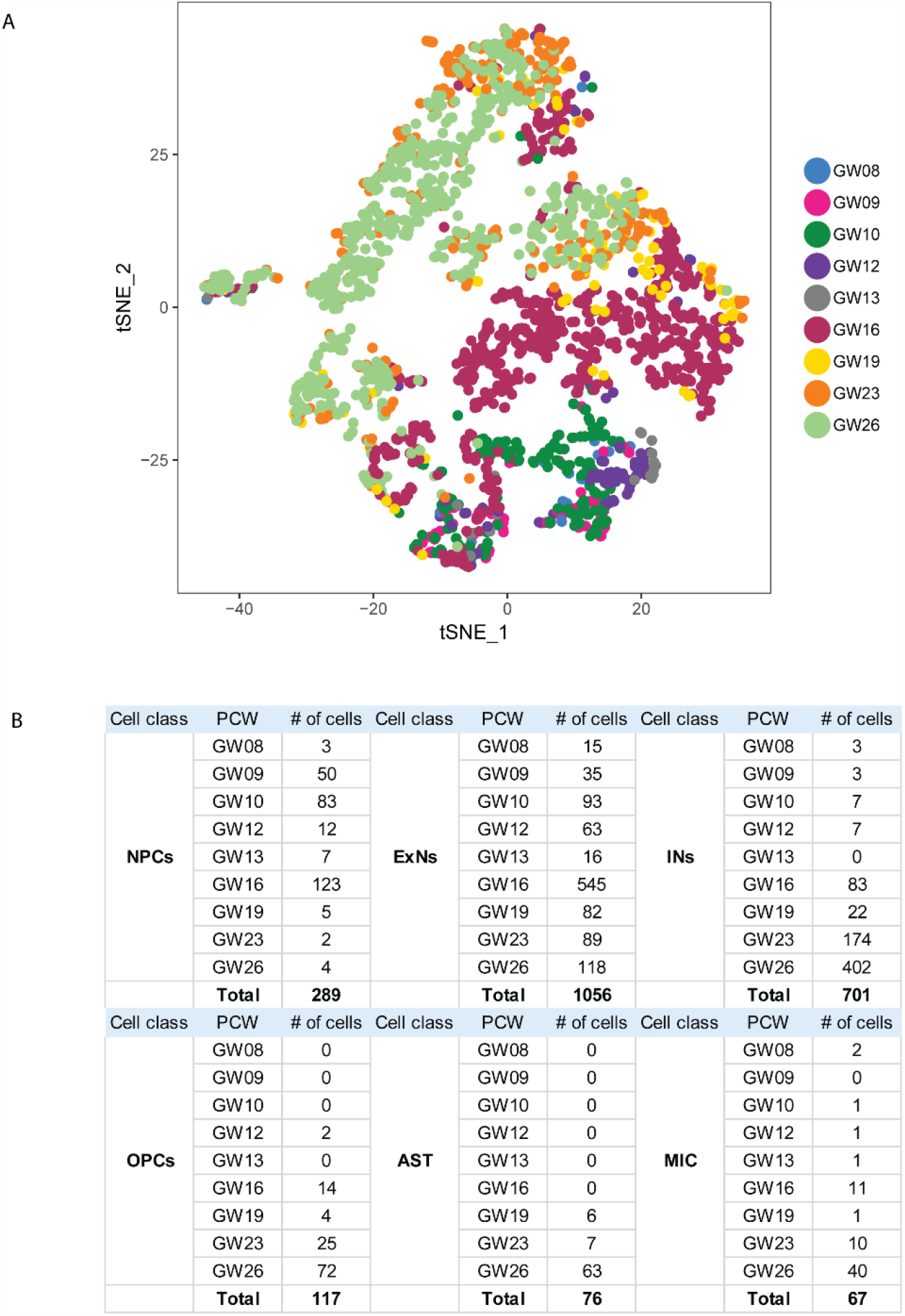
**(A)** *t*-SNE plot showing the distribution of foetal stages **(B)** table showing numbers of cells of each cardinal class at each foetal stage.

**Figure S2.**
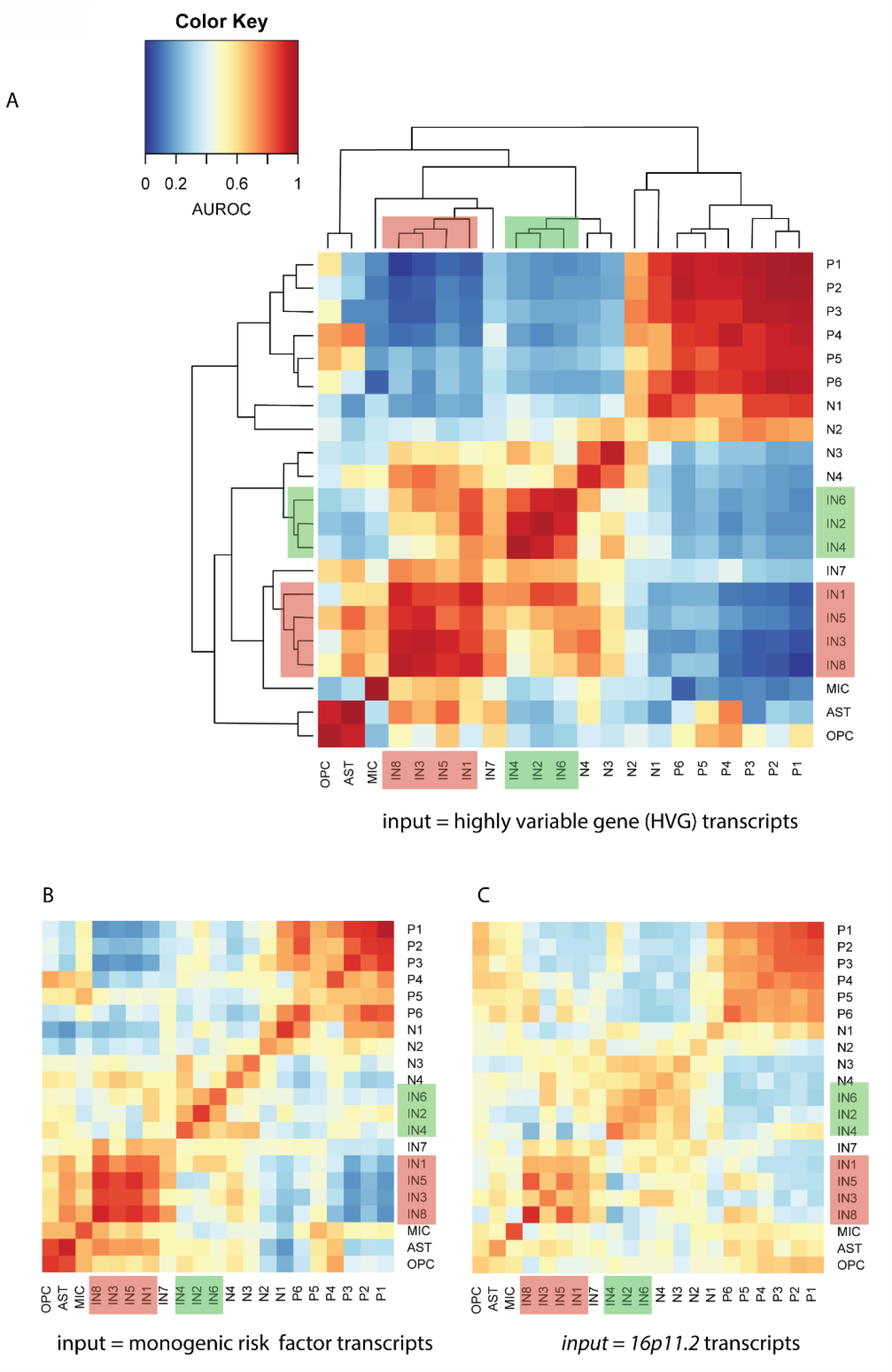
Pairwise comparison of the cluster similarity calculated by MetaNeighbor between the 21 cell clusters. AUROC scores represented as a heatmap where high similarity between clusters is coloured red and low similarity blue. Three plots are shown using different input gene sets **(A)** ∼2000 highly variable gene transcripts between clusters. **(B)** the 83 high confidence and strong candidate (SFARI lists 1 and 2) monogenic autism risk factor transcripts. **(C)** the 27 *16p11.2* transcripts.

**Figure S3.**
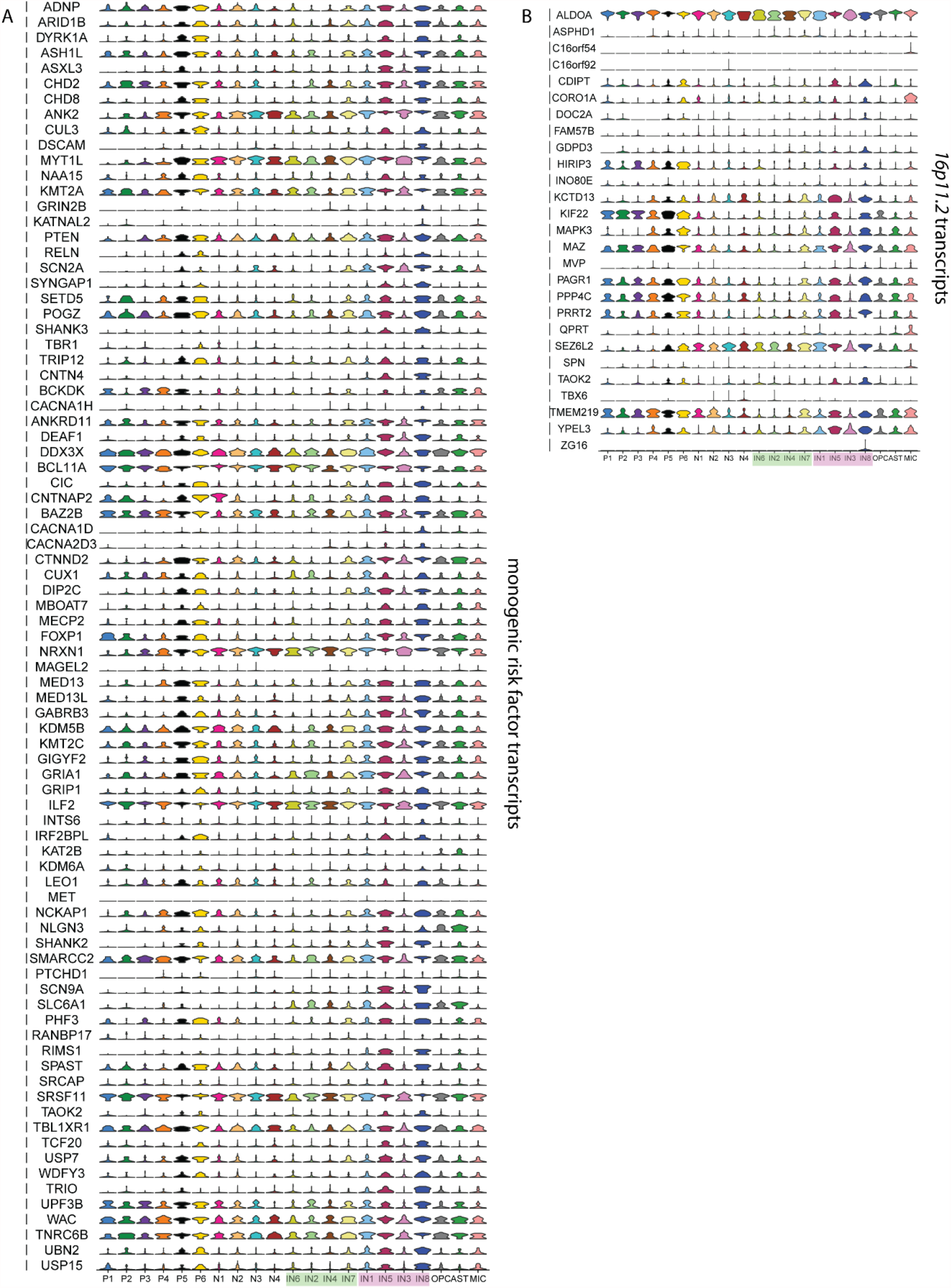
Violin plots showing transcript levels in the 21 different clusters for **(A)** the 83 high confidence and strong candidate (SFARI lists 1 and 2) monogenic autism risk transcripts and **(B)** the 27 *16p11.2* transcripts.

**Figure S4.**
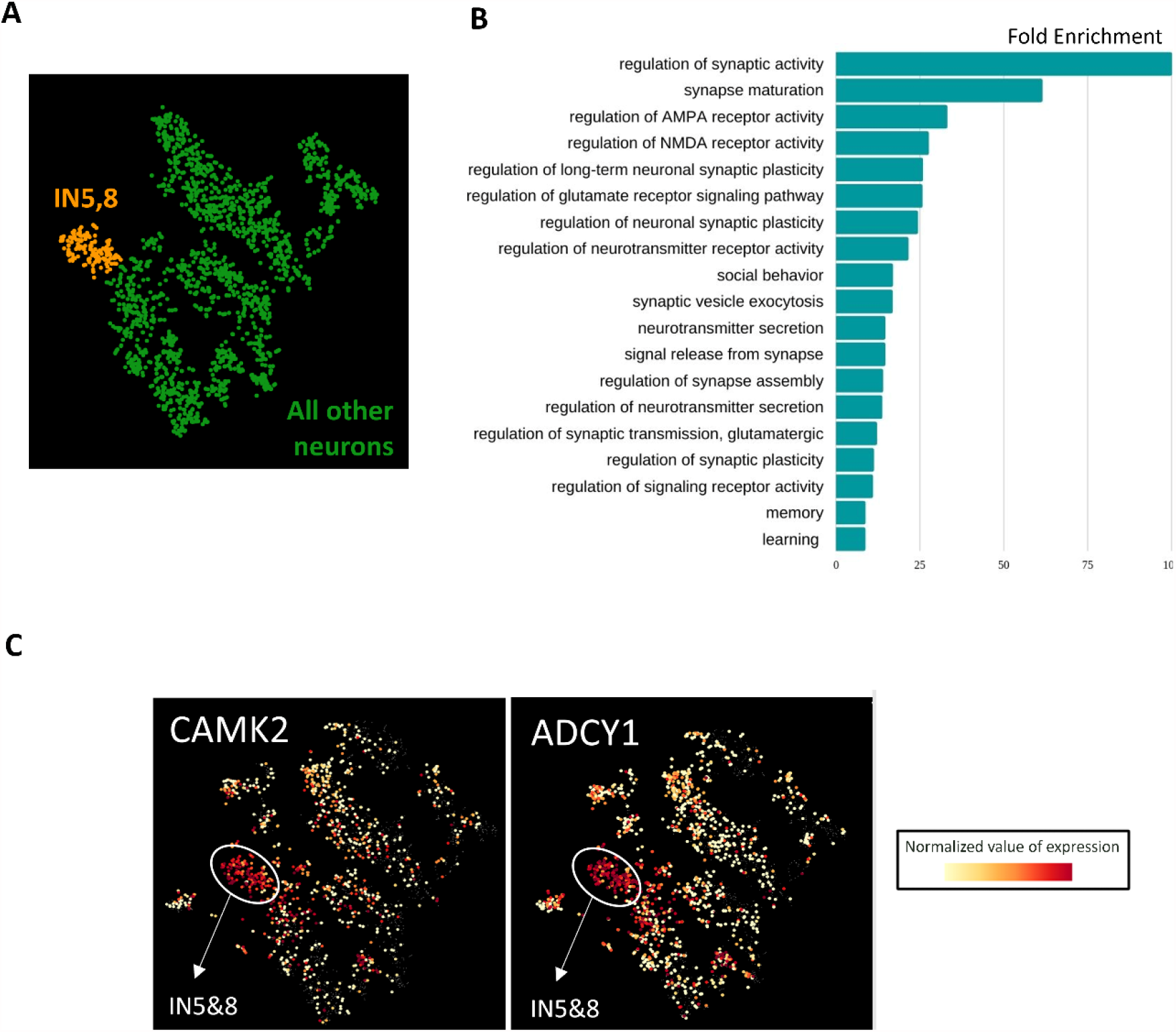
Characterisation of INs by gene ontology analysis. Gene ontology (GO) analysis in IN5,8 (orange in A) versus all other cells (green in A) reveals enrichment of GO terms associated with (B) synaptic activity, maturation, and plasticity. (C) gradient plots of CAMK2, ADCY1 showing that these transcripts are most highly expressed in IN5,8.

**Figure S5:**
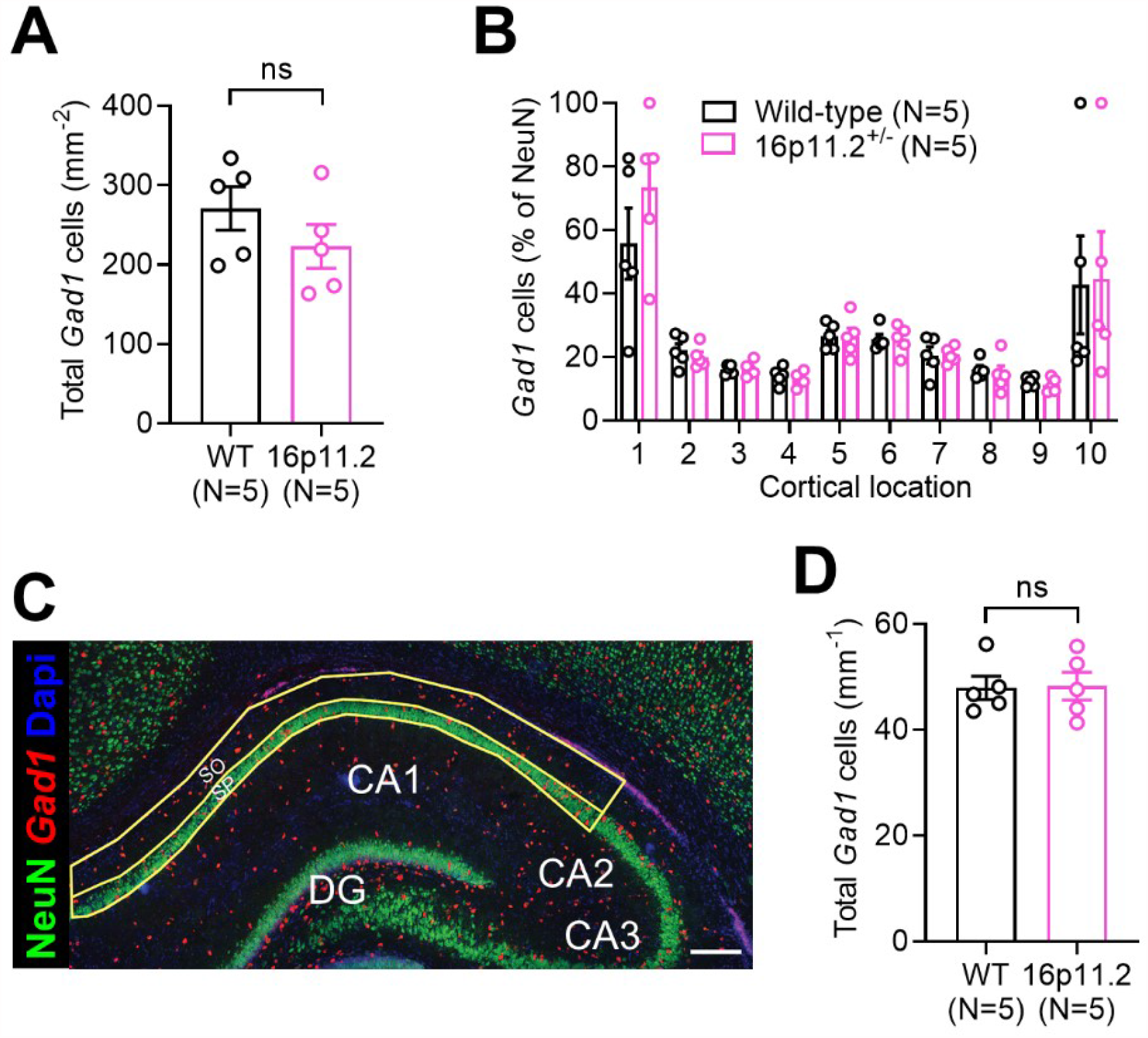
No change in total IN number in somatosensory cortex or hippocampus of the *16p11.2*^+/-^ rat. **(A)** Based on the expression of *Gad1* mRNA, quantification of the total number of INs across the whole somatosensory column from WT (N=6) and *16p11.2*^+/-^ (N=6) rats. (**B**) No change in the relative ratio of *Gad1*-positive cells to total neurons (NeuN) was observed within the cortical column. (**C**) Expanded view of CA1 of the hippocampus from the same image as in Figure 5A, showing Gad1 in situ hybridisation (red), NeuN immunolabelling (green) and DAPI nuclei (blue). Regions used for cell counts in *str. pyramidale* (SP) and *str. oriens* (SO) are delineated with yellow lines. Scale bar: 200 µm. (**D**) Total number of *Gad1*-positive neurons measured in CA1 from WT (N=5) and *16p11.2*^+/-^ (N=5) rats. Statistics shown: ns – p>0.05 from Student’s 2-tailed t-test.

## SUPPLEMENTARY TABLES

**Table S1.**
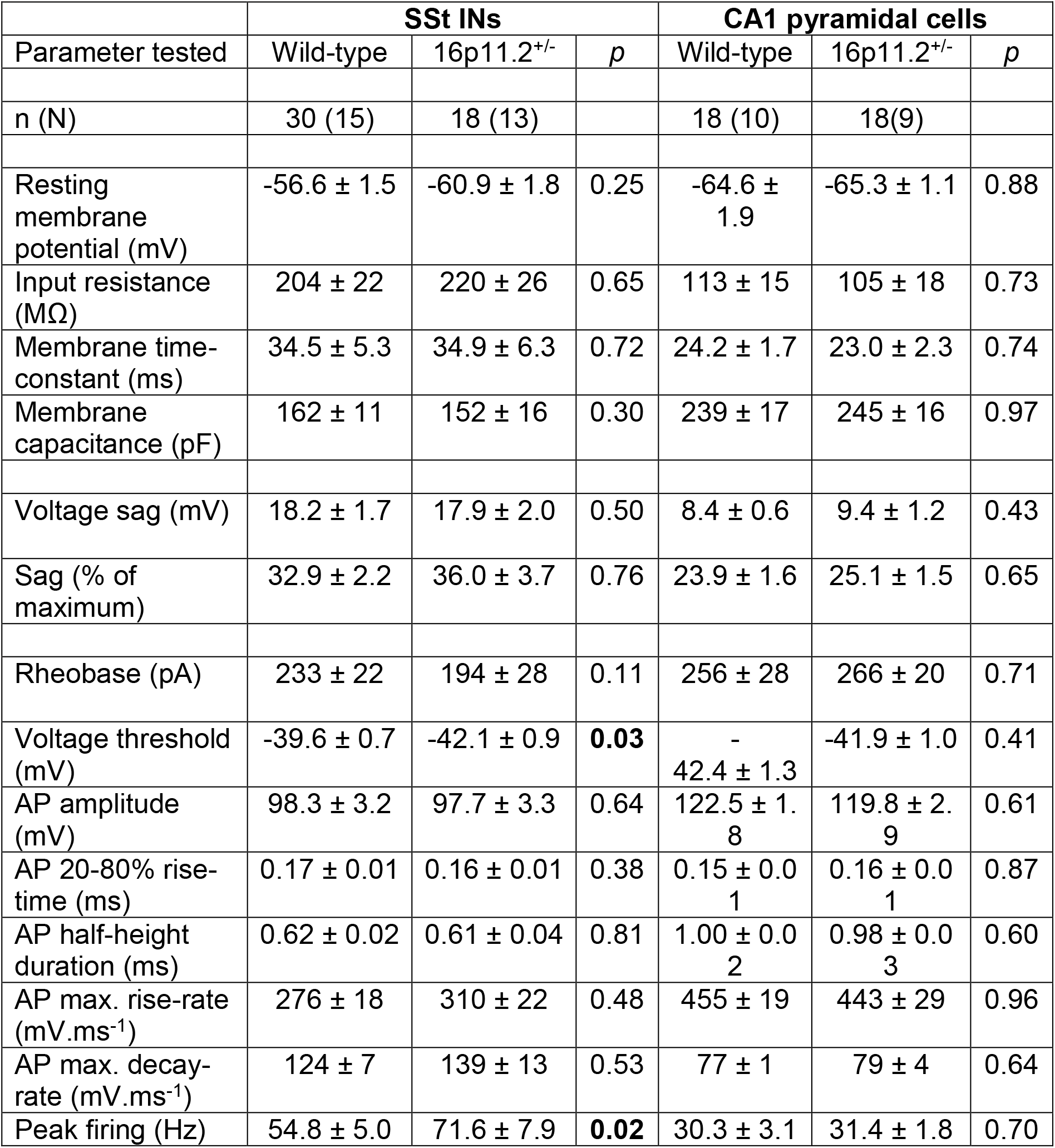

